# Structural analysis of *Sulfolobus solfataricus* TF55β chaperonin in open and filamentous states

**DOI:** 10.1101/2020.01.13.905216

**Authors:** Yi Cheng Zeng, Meghna Sobti, Alastair G Stewart

## Abstract

Chaperonins are biomolecular complexes that assist protein folding. Thermophilic Factor 55 (TF55) is a group II chaperonin found in the archaeal genus *Sulfolobus* that has α, β and γ subunits. Using cryo-electron microscopy, we have determined the structure of the β-only complex of *S. solfataricus* TF55 complexes to 3.6–4.2 Å resolution and a filamentous form to 5.2 Å resolution. The structures of the TF55β complexes formed in the presence of ADP or ATP highlighted an open state in which nucleotide exchange can occur before progressing in the refolding cycle. The structure of the filamentous state indicates how helical protrusions facilitate end-on-end interactions.

**Synopsis:** The isolated complex and filamentous forms of TF55β chaperonin from the thermophilic archaea *Sulfolobus solfataricus* are reported. Using cryo-EM, nucleotide-bound complexes of TF55β at 3.6–4.2 Å resolution reveal an open conformation, while a 5.2 Å reconstruction of the filamentous chaperonin reveals contacts at the apical domain similar to crystal-packed structures.

## 1. Introduction

Chaperonins are large protein complexes that aid in both protein folding and in preventing aggregate-formation by misfolded proteins (Lopez *et al*., 2015; Skjærven *et al*., 2015; Saibil, 2013).

Chaperonins are typically up-regulated during cellular stress or heat-shock to form large (∼1 MDa) molecular cages. These molecular cages are constructed from one or more types of subunits and can capture misfolded protein substrates into their inner cavity. Refolding of the misfolded substrate is achieved by the chaperonin closing around it, which perturbs the local energy landscape of the protein chain. Group I chaperonins, such as GroEL from *E. coli*, form tetradecamers of end-to-end stacked heptameric rings and require heptameric co-chaperones (termed GroES) to close and mediate refolding of the captured protein (Skjærven *et al*., 2015; Weiss *et al*., 2016). Group II chaperonins, such as the eukaryotic TRiC/CCT or archaeal TF55, are similarly composed of two octameric or nonameric stacked rings, but do not require a co-chaperone. However, the group II chaperonins lack an equivalent gene for GroES and instead this function is performed by an additional domain termed the “apical” domain which closes on the central cavity to mediate refolding. The functional and substrate specificity of these chaperonins is mediated by their subunit composition and geometry that work together in a highly cooperative manner (Bigotti & Clarke, 2008; Skjærven *et al*., 2015; Lopez *et al*., 2015) with energy being provided by adenosine triphosphate (ATP) binding and hydrolysis.

Thermophilic Factor 55 (TF55/rosettasome) is a chaperonin found in *Sulfolobus*, a genus of archaea typically residing in volcanic springs that have temperatures around 75–80°C and a pH of 2–3 (Trent *et al*., 1991; Quehenberger *et al*., 2017). As a result of the harsh environment in which *Sulfolobus* live, the TF55 chaperonin is crucial for the survival of these archaea. At temperatures above 80 °C, TF55 is expressed at high levels in *Sulfolobus* due to the presence of a heat-inducible promoter for the *tf55* gene (Jonuscheit *et al*., 2003; Trent *et al*., 1990). However, TF55 regulation and function varies depending on environmental conditions, so that the subunit composition and geometry of the chaperonin is regulated in a temperature-dependent manner, altering between α_6_β_6_γ_6_, α_8_β_8_, or β_18_ complexes, respectively as temperature increases (Chaston *et al*., 2016; Kagawa *et al*., 2003). The ∝, β, and γ subunits share >50 % sequence identity, adopting a similar overall fold, the subunits form either 16-mer or 18-mer complex based on the composition of subunits expressed (Chaston *et al*., 2016; Kagawa *et al*., 2003). The predominant complex formed under heat shock (>80 °C) contains 18 β subunits and is a highly stable and simple system for structural studies (Chaston *et al*., 2016). Filaments of TF55 chaperonin have also been observed in *S. shibatae*, which have been suggested to be a component of the archaeal cytoskeleton (Trent *et al*., 1997, 1998). The ability of the TF55 chaperonin to self-assemble into filaments has the potential to be exploited in the nanobiotechnology field to generate protein nanowires and nanostructures (Li *et al*., 2007; Whitelam *et al*., 2009).

To understand the structure of TF55 in greater detail, we used cryo-electron microscopy (cryo-EM) to image TF55 from the thermophilic archaea *S. solfataricus* in both the single octadecameric complexes and the filamentous form. Complexes composed of only the β subunit (TF55β) were used to aid in purification and homogeneity of the sample. Reconstructions of TF55β complexes in the presence of either ATP or ADP resulted in similar open structures with an accessible binding site. Filaments of TF55β were observed at high protein concentrations (> 10 mg/mL) or could be induced by the addition of dodecyl maltoside (DDM). Three-dimensional (3D) reconstructions of these filaments showed that polymerization is mediated by the helical protrusions.

## 2. Materials and methods

### 2.1. Purification of TF55β complex from *S. solfataricus*

The TF55β complex was purified from *S. solfataricus* as described previously (Chaston *et al*., 2016). *S. solfataricus* (DSM1617) cell paste was purchased from the University of Georgia Biofermentation Facility. Cells were grown at 80 °C (pH 3.7) and harvested at an OD_600_ of 1.2. 20 g of cell paste was resuspended in 350 mL of lysis buffer (50 mM Tris/Cl pH 8.0, 5 mM MgCl_2_, 0.001% PMSF, 100 mM NaCl, 1% DDM) and sonicated using a Branson Digital Sonifier for 3 min at 30 % amplitude with 1 s on/off cycles. The lysate was centrifuged in a Beckman 45Ti rotor at 100,000 x g for 30 min at 4 °C and the supernatant applied to a HiLoad 26/10 Q Sepharose column (GE Life Sciences) equilibrated in 90 % buffer A and 10% buffer B (buffer A: 20 mM Tris/Cl pH 8.0, 2 mM MgCl_2_, 1 mM EDTA, 0.001% PMSF, 0.05% DDM; buffer B: same as A with 1 M NaCl). The column was washed until a constant Abs_280nm_ baseline was established after which the protein was eluted in a 10-50% buffer B gradient over 400 mL. Fractions containing TF55 were pooled and concentrated to less than 10 mL before applying to a HiPrep 16/60 Sephacryl S-300 column equilibrated in buffer C (20 mM Tris/Cl pH 8.0, 2 mM MgCl_2_, 1 mM EDTA, 100 mM NaCl). Fractions corresponding to the β subunit of TF55 (as assessed by mass spectrometry finger printing) were pooled and Mg•ATP or Mg•ADP was added to a final concentration of 2 mM. Each sample was concentrated to 0.5 mL and applied to a Superose 6 10/300 GL column equilibrated in buffer C to obtain the TF55β complex in the presence of ATP or ADP (Supplementary Figure S1a,b). The protein was concentrated to either 20.0 mg/mL (ATP-complex) or 24.0 mg/mL (ADP-complex), which was then flash frozen and stored at –80 °C before use.

### 2.2. Cryo-EM data collection and preprocessing for single particle TF55β complex

A 3.5 µL droplet of purified TF55β complex at either 20.0 mg/mL (ATP-complex) or 12.0 mg/mL (ADP-complex) was applied to a copper grid with holey carbon film (Quantifoil Copper R1.2/1.3, 200 mesh) that had been glow discharged for 60 s. Grids were blotted for 5 s at 20°C and flash-frozen in liquid ethane using a FEI Vitrobot Mark IV.

Grids were transferred to a FEI Talos Arctica transmission electron microscope operating at 200 kV. Images were recorded automatically using EPU at a magnification of 150,000 ×, yielding a pixel size of 0.986 Å. A total dose of 50 electrons (spread over 29 frames) per Å^2^ and a total exposure time of 61 s were used on the Falcon-III detector operating in counting mode. A total of 1641 movies (ATP-complex) or 2321 movies (ADP-complex) were collected.

The data processing workflow for single particle analysis of the ATP- and ADP-complex is summarized in Supplementary Figure S1d. Movies were imported into cryoSPARC2 (Punjani *et al*., 2017) and the built-in patch motion correction program was used to correct for local beam-induced motion and to align the frames. Defocus and astigmatism values were estimated using the built-in patch CTF estimation program. Micrographs were excluded on the basis of drift, excessive ice contamination, or excessive filament formation, resulting in 1632 (ATP-complex) or 1463 (ADP-complex) micrographs.

### 2.3. Data processing and refinement for single particle TF55β complex

Different picking strategies were required for the two datasets. For the ATP-complex dataset, an initial 28 micrographs were manually picked for side-view particles (Supplementary Figure S1d) to generate a 2D template which was used to pick a separate 19 micrographs, resulting in 1521 particles that encapsulated multiple orientations of the complex. The extracted particles (with a box size of 320^2^ pixels resized to 160^2^ pixels) were used as the training data for the neural network-based picker Topaz (Bepler *et al*., 2019). Particles were down-sampled by a factor of 16 with 200 expected particles per micrograph. Topaz picked a total of 213,286 particles across all 1632 micrographs. For the ADP-complex dataset, the built-in blob picker in cryoSPARC2 was used to pick a total of 395,079 particles across all 1463 micrographs.

For both datasets, particles were extracted at a box size of 440^2^ pixels and subject to iterative rounds of 2D classification. Side views of the particles were rigorously separated from filamentous segments present in the 2D class average by iterative classification with the maximum alignment resolution set at 20 Å. A total of 94,618 (ATP-complex) or 133,319 particles (ADP-complex) were obtained after 2D classification. 3D *ab initio* reconstruction with two classes set to an expected similarity of zero and no symmetry set (C_1_) was used as a surrogate for 3D classification, separating top views and “junk” particles from particles that contributed towards the final reconstruction of the complex. No other forms or conformations of TF55β (such as the closed state) in both datasets could be found when the number of 3D classes was increased beyond two. 3D classification resulted in a final 53,777 (ATP-complex) or 43,528 (ADP-complex) particles carried forward to 3D refinement. Homogenous refinement in C_1_ gave maps with an estimated resolution of 4.19 Å (ATP-complex) or 6.08 Å (ADP-complex) based on the gold-standard Fourier Shell Correlation (FSC = 0.143) criterion. Local motion correction and subsequent homogenous and non-uniform refinement with D_9_ symmetry imposed generated maps with an estimated resolution of 3.62 Å (ATP-complex) and 4.19 Å (ADP-complex). Local resolution was calculated using the implementation in cryoSPARC2. Refinement was also attempted in Relion2 (Zivanov *et al*., 2018), however it did not classify a closed-state chaperonin or produce a better map compared to cryoSPARC2. Cryo-EM collection and refinement statistics are listed in Table 1.

**Table 1.**
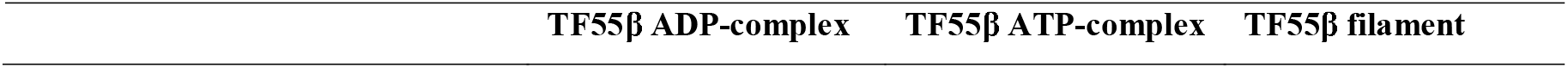
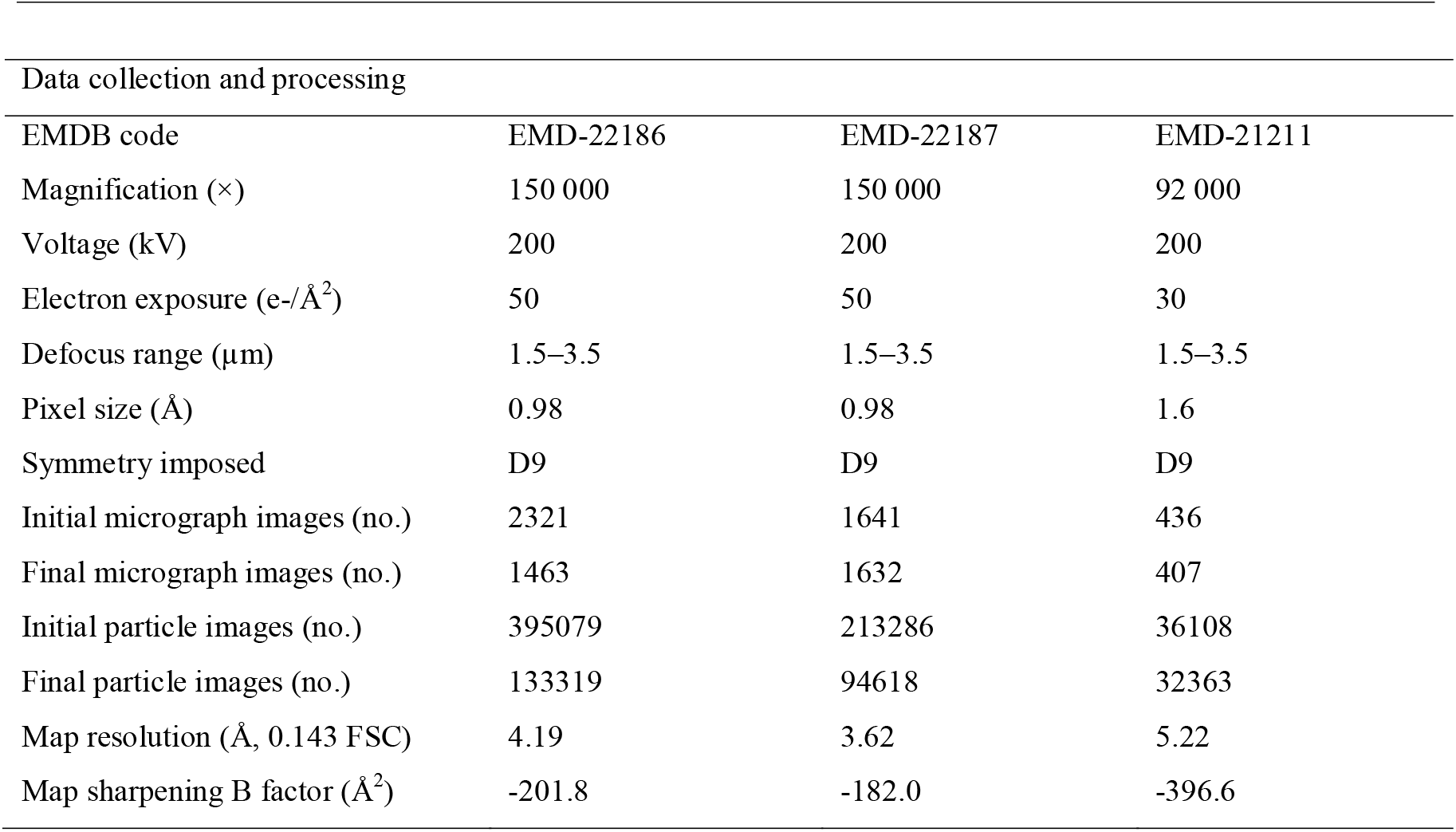
Data-collection and coordinate model-refinement statistics

| TF55β ADP-complex | TF55β ATP-complex | TF55β filament |
| --- | --- | --- |

| Data collection and processing |  |  |  |
| --- | --- | --- | --- |
| EMDB code | EMD-22186 | EMD-22187 | EMD-21211 |
| Magnification (×) | 150 000 | 150 000 | 92 000 |
| Voltage (kV) | 200 | 200 | 200 |
| Electron exposure (e-/Å<sup>2</sup>) | 50 | 50 | 30 |
| Defocus range (µm) | 1.5–3.5 | 1.5–3.5 | 1.5–3.5 |
| Pixel size (Å) | 0.98 | 0.98 | 1.6 |
| Symmetry imposed | D9 | D9 | D9 |
| Initial micrograph images (no.) | 2321 | 1641 | 436 |
| Final micrograph images (no.) | 1463 | 1632 | 407 |
| Initial particle images (no.) | 395079 | 213286 | 36108 |
| Final particle images (no.) | 133319 | 94618 | 32363 |
| Map resolution (Å, 0.143 FSC) | 4.19 | 3.62 | 5.22 |
| Map sharpening B factor (Å<sup>2</sup>) | -201.8 | -182.0 | -396.6 |

For model building, chain A of the crystal structure of TF55β (PDB ID: 4XCD, Chaston et al., 2016) was docked as a rigid body into the density of one of the subunits in the D_9_ symmetric cryo-EM map using Chimera’s fitting program (Pintilie *et al*., 2010). Modelling was performed using Coot (Emsley *et al*., 2010), Phenix real-space refinement (Adams *et al*., 2010), and ISOLDE (Croll, 2018). For the apical domain (residues 203-381), molecular dynamics flexible fitting (MDFF) in ISOLDE with secondary structure restraints allowed for modelling of major β-sheet and α-helical densities present at lower resolutions. Density for loops in the apical domain was not sufficient to enable them to be modelled accurately and so these were omitted from the coordinates that were deposited to the Protein Data Bank. Model building and refinement statistics are listed in Table 2. Figures were prepared using PyMOL (Schrodinger, LLC), Chimera (Pettersen *et al*., 2004) or ChimeraX (Goddard *et al*., 2018).

**Table 2.**
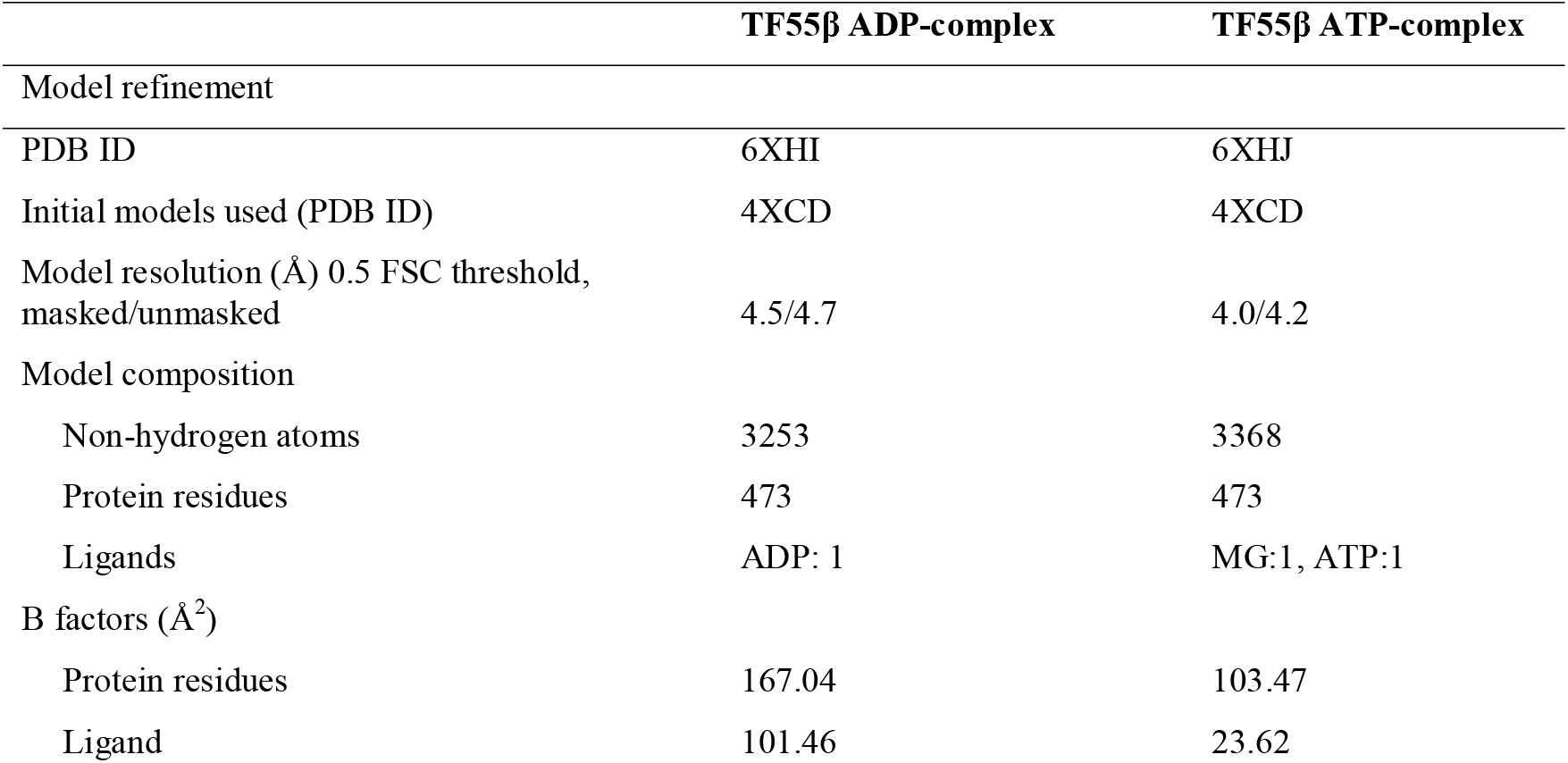
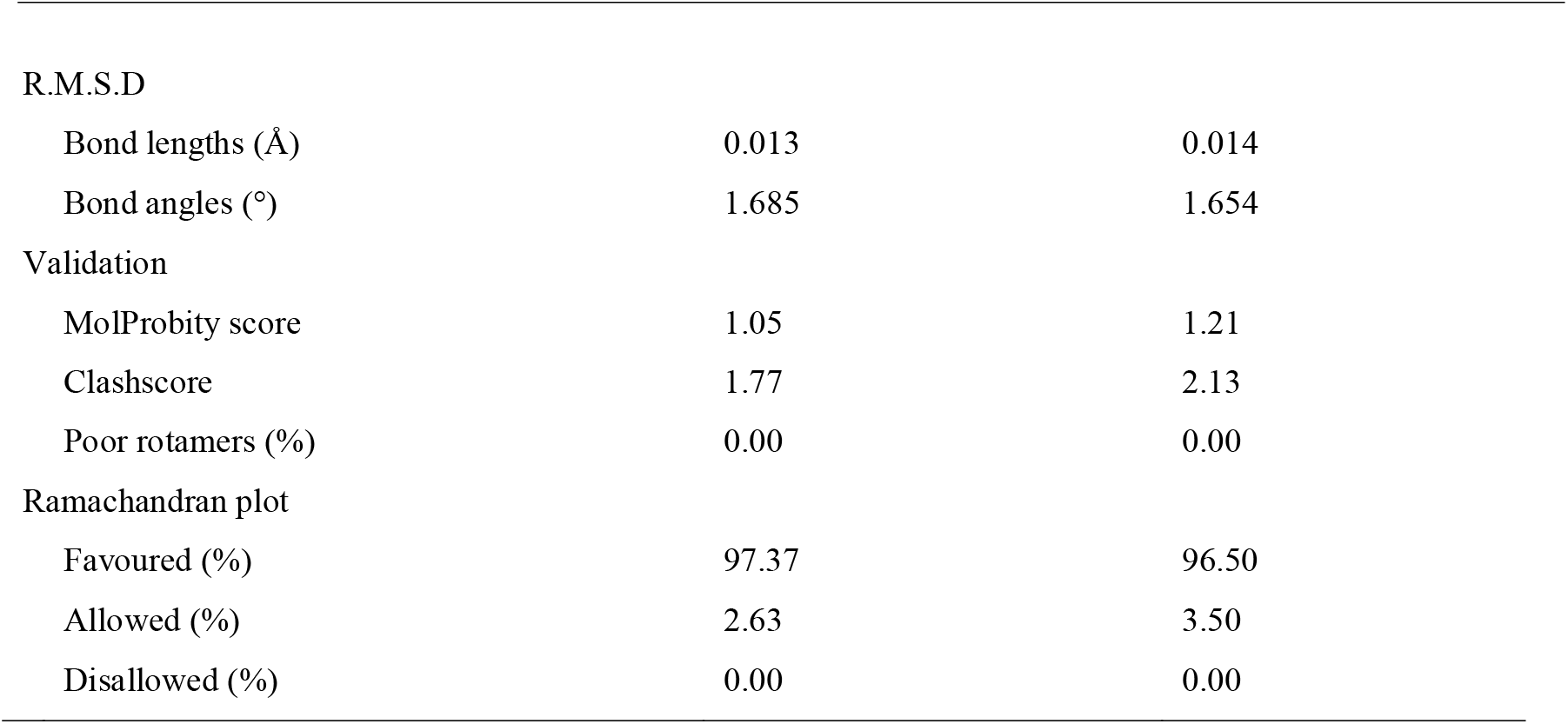
Coordinate model-refinement statistics

|  | TF55β ADP-complex | TF55β ATP-complex |
| --- | --- | --- |
| Model refinement |  |  |
| PDB ID | 6XHI | 6XHJ |
| Initial models used (PDB ID) | 4XCD | 4XCD |
| Model resolution (Å) 0.5 FSC threshold, masked/unmasked | 4.5/4.7 | 4.0/4.2 |
| Model composition |  |  |
| Non-hydrogen atoms | 3253 | 3368 |
| Protein residues | 473 | 473 |
| Ligands | ADP: 1 | MG:1, ATP:1 |
| B factors (Å<sup>2</sup>) |  |  |
| Protein residues | 167.04 | 103.47 |
| Ligand | 101.46 | 23.62 |
| R.M.S.D |  |  |
| Bond lengths (Å) | 0.013 | 0.014 |
| Bond angles (°) | 1.685 | 1.654 |
| Validation |  |  |
| MolProbity score | 1.05 | 1.21 |
| Clashscore | 1.77 | 2.13 |
| Poor rotamers (%) | 0.00 | 0.00 |
| Ramachandran plot |  |  |
| Favoured (%) | 97.37 | 96.50 |
| Allowed (%) | 2.63 | 3.50 |
| Disallowed (%) | 0.00 | 0.00 |

### 2.4. Cryo-EM data collection and preprocessing for filamentous TF55β

TF55β chaperonin (ATP-complex) was diluted with buffer C containing 0.01% DDM to obtain a final protein concentration of 10 mg/ml and DDM concentration of 0.005%, and incubated on ice for 30 min. A 3.5 µL droplet of the sample was applied to a copper grid with holey carbon film (Quantifoil Copper R1.2/1.3, 200 mesh) that had been glow discharged for 60 s. Grids were blotted for 4 s at 20 °C and flash-frozen in liquid ethane using a FEI Vitrobot Mark IV.

Grids were transferred to a FEI Talos Arctica transmission electron microscope operating at 200 kV. Images were recorded automatically using EPU at a magnification of 92,000 ×, yielding a pixel size of 1.6 Å. A total dose rate of 30 electrons (spread over 30 frames) per Å^2^ per second and a total exposure time of 116 s were used on the Falcon-III detector operating in counting mode. 436 movies were collected in total. The data processing workflow for the TF55β filament is summarized in Supplementary Figure S4a. Micrographs were imported into cryoSPARC2 (Punjani *et al*., 2017) and motion correction and CTF estimation were performed using the Patch motion or Patch CTF programs respectively. 29 micrographs were discarded on the basis of drift or excessive ice contamination.

### 2.5. Data processing and refinement for filamentous TF55β

From a 2D template of filaments that were manually traced using e2helixboxer.py (Tang *et al*., 2007) and 2D classified, segments of filaments were automatically picked in cryoSPARC2 using the template picker to obtain 36,108 segments from 406 micrographs. After two rounds of 2D classification to filter out junk particles and bad segments, a final 32,363 segments with a box size of 300 pixels were then subjected to *ab initio* reconstruction with D_9_ symmetry applied in order to obtain a segment with two full chaperonins. Further homogenous and non-uniform refinement after re-extracting particles from local motion correction converged on a map with an estimated resolution of 5.22 Å based on the gold-standard Fourier Shell Correlation (FSC = 0.143) criterion. Docking and manual fitting was initially performed in ChimeraX/ISOLDE using two chains of the cryo-EM model of TF55β chaperonin orientated at the helical protrusion interface. The resulting coordinates were used as the starting model for further MDFF in NAMD (Trabuco *et al*., 2008).

### 2.6. Filament formation screening

TF55β was purified either without any nucleotide, Mg•ATP or Mg•ADP and concentrated to 3 mg/mL, 9 mg/mL or 12 mg/mL, respectively. Concentrated protein was either directly used for grid preparation or further incubated with DDM (0.005%), AlF_x_ (300 mM NaF, 50 mM Al(NO_3_)_3_.9H_2_O), ATP.AlF_x_ (10 mM ATP, 300 mM NaF, 50 mM Al(NO_3_)_3_.9H_2_O), and/or ADP.AlF_x_ (10 mM ADP, 300 mM NaF, 50 mM Al(NO_3_)_3_.9H_2_O) as shown in Supplementary Figure S3. A 3.5 µL droplet of the sample was applied on a copper grid with holey carbon film (Quantifoil Copper R1.2/1.3, 200 mesh) that had been glow discharged for 60 s. Grids were blotted for 4 s and flash-frozen in liquid ethane using a FEI Vitrobot Mark IV. Grids were transferred to a FEI Talos Arctica transmission electron microscope operating at 200 kV. Images were recorded automatically using EPU at a magnification of either 92,000 × or 150,000 × with a total dose of 40 electrons per Å^2^ per second.

## 3. Results and discussion

### 3.1. Single particle analysis of the TF55β complex

Cryo-EM micrographs of TF55β chaperonin formed in the presence of either ATP or ADP showed nonameric ring particles (Figure 1). 3D reconstruction of these particles showed both ATP- and ADP-bound octadecameric complexes in an open configuration and had resolution estimates of 3.6 Å and 4.2 Å, respectively (Figure 1 and Supplementary Figure S1). Attempts to 3D classify particles into different lid conformations did not yield any discrete classes (data not shown). Sidechain density for the equatorial and intermediate domain was more evident than the apical domain, where only the mainchain of the secondary structures could be modelled (Supplementary Figure S1). Clearer density for the loops in the equatorial domain were observed in the cryo-EM map compared to the electron density map of the previously solved crystal structure used as the template (PDB ID: 4XCD). Because the map density was weak for the apical domain, modelling was performed using molecular dynamics flexible fitting (MDFF) with secondary structure restraints.

**Figure 1.**
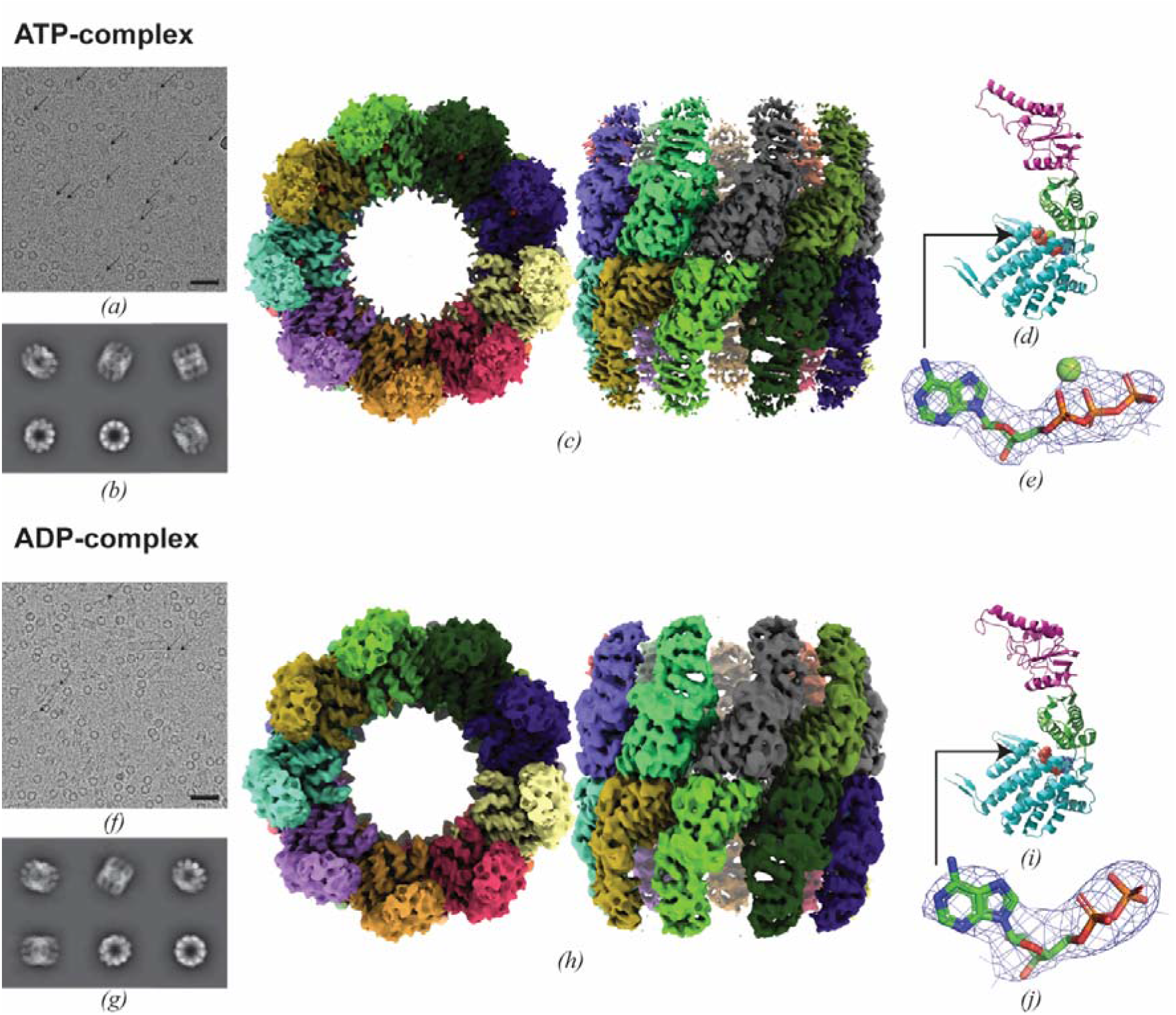
The cryo-EM structure of the octadecameric TF55β complex with either ATP or ADP bound. *(a, f)* Representative cryomicrograph of TF55β chaperonins. Arrows indicate side and tilted views of the complex. Scale bars are 50 nm. *(b, g)* Selected 2D class averages of TF55β chaperonin from particles contributing to the final 3D reconstruction map. Cryo-EM map of TF55β chaperonin viewed from the top and side *(c, h)*, with colors segmenting each corresponding subunit of the homo-octadecamer. *(d, i)* Cartoon representation of a subunit of the TF55β complex, with the domains colored and labelled (apical, magenta; intermediate, green; equatorial, cyan). Cryo-EM density (blue mesh) around Mg•ATP contoured at σ = 7.0 *(e)* and ADP contoured at σ = 9.0 *(j)* within the binding site of a subunit of the TF55β complex.

There was very little variability between the two models that showed an overall RMSD of 1.0 Å and had only minor changes around the nucleotide binding site (Figure 2). In contrast with the crystal structure solved previously (Chaston *et al*., 2016), the apical domain was hinged at the helices of the intermediate domain and this generated an outward tilt in the EM structures (Figure 2a), with the aperture into the central cavity increased. Our cryo-EM model likely reflects the structure of the chaperonin in solution, devoid of crystal contacts between complexes that may induce crystal packing artefacts. The highly flexible apical domain plays a key role in the opening/closing of the complex to facilitate substrate binding and refolding, and this flexibility is conserved throughout chaperonins in other organisms (Chaudhry *et al*., 2004; Zang *et al*., 2016). The lower resolution achieved in the intermediate and apical domains (Supplementary Figure S1) probably indicates that these regions are more dynamic in solution.

**Figure 2.**
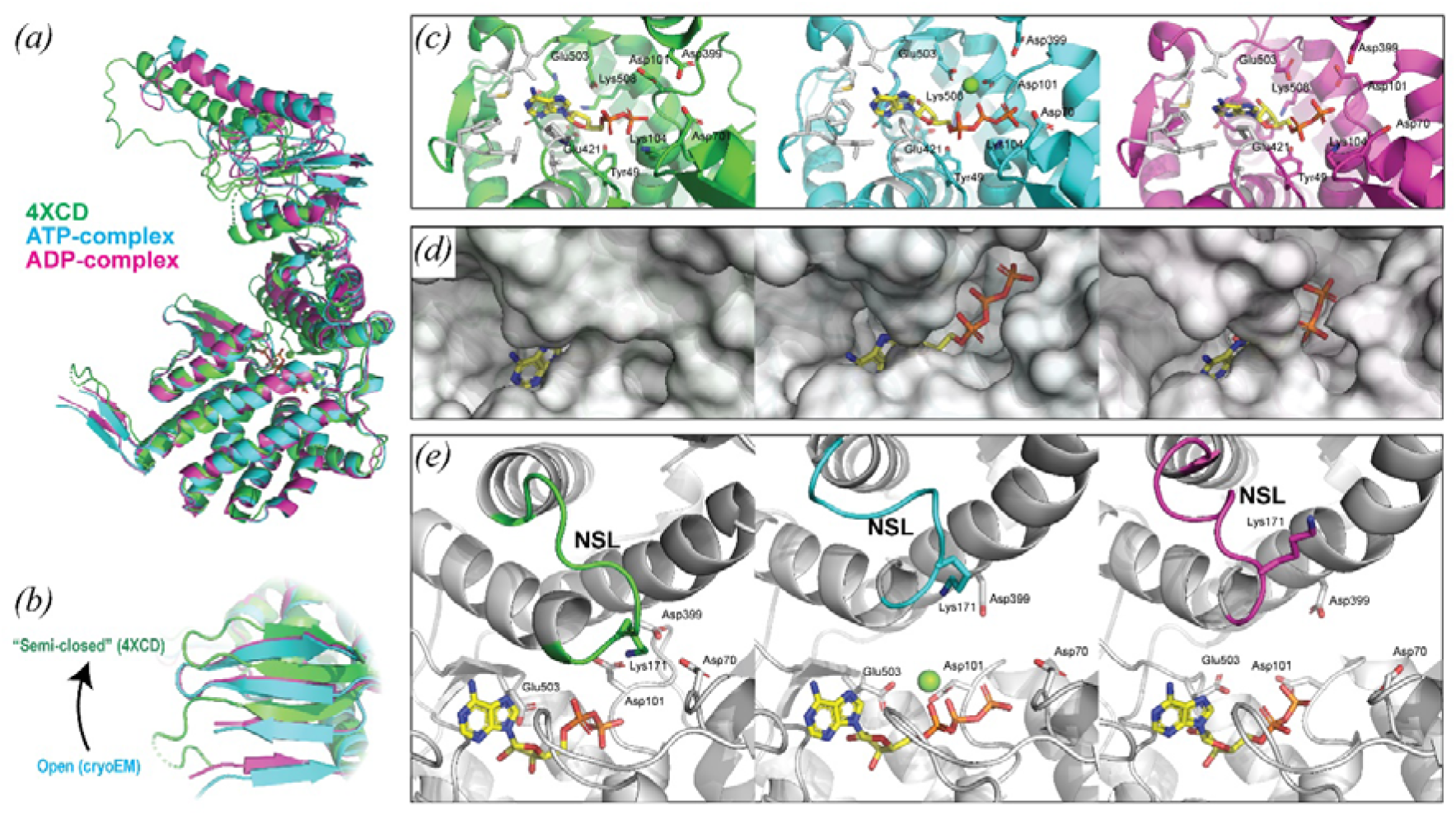
Comparison of the TF55β crystal structure with the open-state TF55β cryo-EM structures. 4XCD crystal structure in green, cryo-EM model of ATP-complex in cyan, cryo-EM model of ADP-complex in magenta. *(a)* Overlay of the atomic models of the crystal structure against cryo-EM models aligned to the equatorial domain showing an inward tilt of the apical and intermediate domains. *(b)* Extended β-sheet of the stem-loop with the C- and N-termini of the adjacent subunit (N-termini not modelled in 4XCD) shifts upwards in the crystal structure (resembling transitory “semi-closed” state between open and closed) relative to the equatorial domain. *(c)* Nucleotide binding site and associated residues. Stick models for hydrophobic residues (in white) and polar residues labelled. *(d)* Protein surface representation shows the nucleotide binding pocket is more accessible in the cryo-EM open state. *(e)* Nucleotide sensing loop (NSL, colored) shifts up in the cryo-EM models, moving Lys171 away from the binding site.

The stem-loop forms an extended β-sheet with the terminal β-strands of the adjacent subunit near the nucleotide binding site connecting adjacent subunits (Figure 2b), although these features are shifted towards the apical domain in the cryo-EM structures relative to the crystal structures of TF55β (Supplementary Figure S2a). Similar changes were observed in crystal structures of open- and closed-state of the group I chaperonin GroEL (Saibil *et al*., 2013) and a group II-like chaperonin from the bacteria *Carboxydothermus hydrogenoformans* (An *et al*., 2017). The stem-loop that forms β-sheets with the N- and C-terminus of the neighboring subunit may coordinate ring closing allosterically through contacts at the base of the loop with the nucleotide, together with conformational changes generated by the binding of a misfolded protein substrate at the equatorial-intermediate domains, as seen in the eukaryotic CCT:tubulin-bound complex (Saibil *et al*., 2013; Muñoz *et al*., 2011).

### 3.2. The open conformation of TF55β has an accessible nucleotide site

Inspection of the nucleotide binding sites in each of the cryo-EM maps clearly indicated the presence of either ATP or ADP for the respective complexes (Figure 1). A pocket of hydrophobic residues surrounds the aromatic base while multiple polar residues coordinate to the ribose sugar and phosphates (Figure 2c). Density present for a magnesium ion in the ATP bound complex shows that it is coordinated to the phosphates of ATP together with Asp101 and Glu503 of TF55β. However, in the intermediate domain in the fully open state it is further away from the catalytic Asp399 (Supplementary Figure S2b) compared to closed state structures (Lopez *et al*., 2015; Pereira *et al*., 2012).

The nucleotide binding pocket in the open cryo-EM structures becomes more solvent accessible compared to that seen in the crystal structure of the same molecule (Figure 2d). The nucleotide sensing loop moves towards the apical domain, with the conserved Lys171 (Pereira *et al*., 2012) flipped away from the nucleotide (Figure 2e). This probably reflects a state where, post-hydrolysis and substrate refolding, ADP is exchanged for ATP and a new folding cycle commences. Interestingly, no significant change in the tilt of the intermediate or apical domain was observed between the ATP-complex and ADP-complex, suggesting that ATP-binding may not be sufficient to induce a conformational change in TF55β. This is in contrast with other group II chaperonins such as that from *M. maripaludis* where a conformational change in the apical domain can be seen upon ATP binding (Zhang *et al*., 2011) or from group I chaperonins such as GroEL where ATP binding causes a tilt in the apical and intermediate domains to facilitate substrate binding and GroES docking (Clare *et al*., 2012). Instead, ATP hydrolysis is likely the source of energy from which conformational changes occurs, leading to a transition to the closed state (Sekiguchi *et al*., 2013). The open and closed EM structures of *A. tengchongensis* recapitulates these findings where incubation with the non-hydrolysable ATP-AlF_x_ prevents lid closing, while the apo structure (after incubation with ATP over 2 weeks) results in only closed chaperonin (Huo *et al*., 2010).

### 3.3. TF55β filaments form at high protein concentrations, but are disassembled by EDTA

At concentrations of 9 mg/ml, short assemblies of TF55β could be observed on grids, and were localized preferentially near the edge of the holes of the holey carbon grid (Figure 3a). At a higher concentration of 20 mg/ml, longer filaments could be observed alongside top views of the TF55β complex (Figure 3b). It also appeared that there was no preference for ADP or ATP (or ATP analogues such as ADP.AlF_x_) to induce filament formation because, in each case, filaments were observed when TF55β concentration was increased (Supplementary Figure S3). The addition of EDTA in excess of the magnesium ion concentration present caused the filaments to dissociate (Figure 3c) as previously observed in *S. shibatae* TF55 (Trent *et al*., 1997), but also visibly reduced the number of TF55β complexes. EDTA is able to release nucleotides from chaperonin complexes (Lin *et al*., 2013) and so TF55 from *S. solfataricus* may be less stable without the presence of both Mg^2+^ and nucleotide (Trent *et al*., 1997). It is uncertain if these filaments are present *in vivo* or are an artefact of sample or grid preparation. Initial attempts to analyze these filaments were hindered by the large number of single particles that were also present which interfered with helical reconstruction.

**Figure 3.**
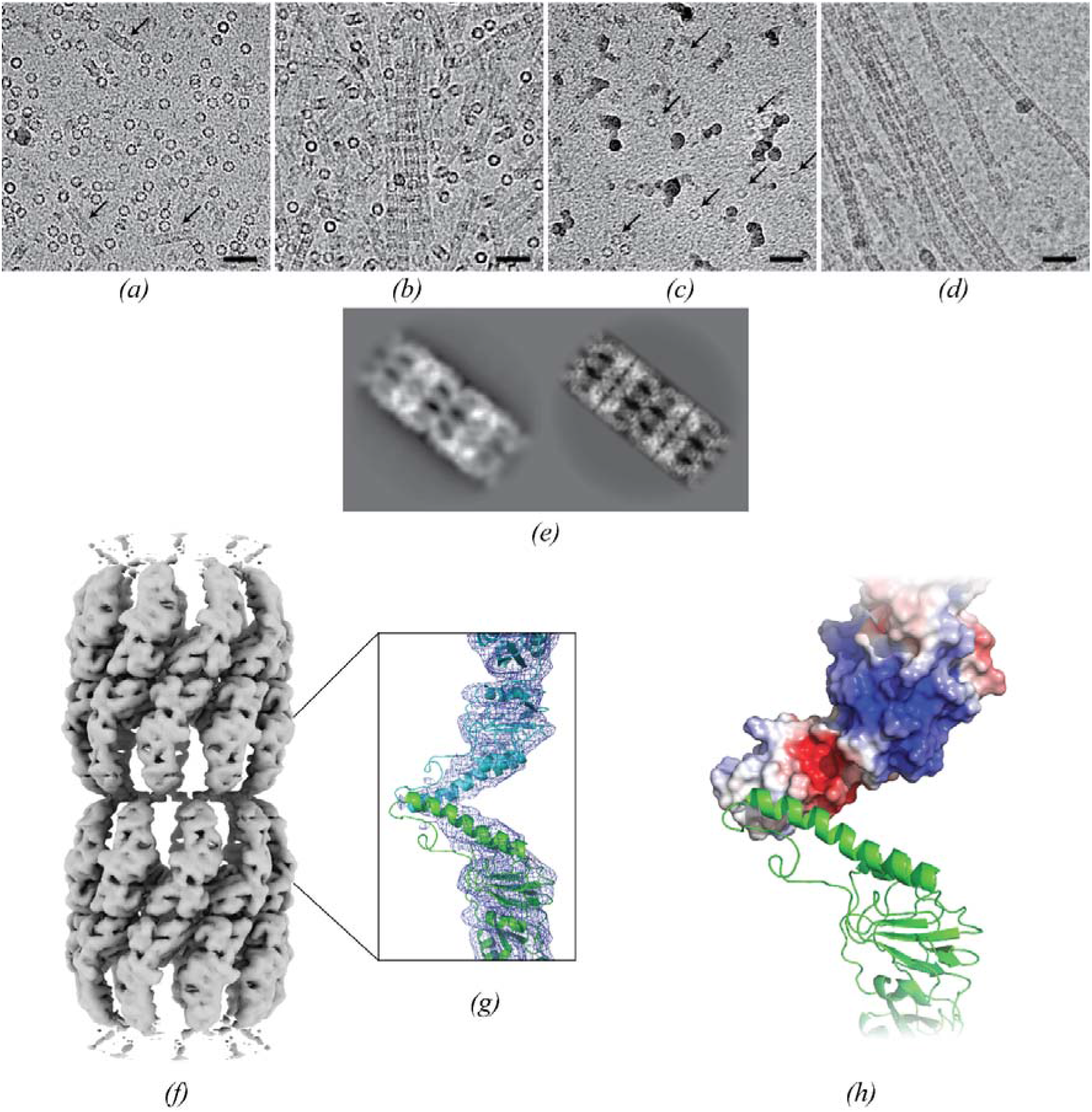
TF55β filament formation. *(a-d)* Micrographs of TF55β chaperonin under different conditions near the edge of the grid holes.Scale bars are 50 nm. *(a)* TF55β at 9 mg/ml, arrows pointed at small assemblies of chaperonins. *(b)* TF55β at 20 mg/ml, showing striated filaments and top views of single complexes (rings). *(c)* TF55β at 19 mg/ml and 5 mM EDTA, showing loss of filaments to single complexes (arrow, ring particles). *(d)* TF55β at 9 mg/ml and 0.005% DDM, showing striated filaments only. *(e)* 2D class average of a segment of the filamentous TF55β (left) and a 2D projection from the crystal lattice of TF55β (right, PDB ID: 4XCD) showing striking similarities in the mode of interaction between complexes. *(f)* Cryo-EM map of a two-chaperonin segment of the filamentous TF55β. *(g)* MDFF of two subunits from the isolated complex model (cartoon) into the cryo-EM map (blue mesh) indicated an interaction near the apex of the helical protrusion. *(h)* Electrostatic surface shows a hydrophobic patch at the end of the protruding helices where the contacts in filament are found. Blue = negative potential, red = positive potential, white = neutral/hydrophobic.

### 3.4. Filamentous TF55β assembly can be induced by DDM

While attempting to overcome the preferential orientation problem common to cryo-EM of group II chaperonins (Zang *et al*., 2016), we found serendipitously that the inclusion of 0.005% DDM could cause the TF55β complex to form long filamentous assemblies without any trace of single particles (Figure 3d), even at a lower concentration of 10 mg/ml. The filaments of TF55β generated with DDM resembled those seen with high protein concentration (Figure 3b) as well as those previously described in *S. shibatae* (Trent *et al*., 1997), showing characteristic striations both at the nonameric double ring of individual TF55β complexes and at the contacts between end-to-end stacked chaperonins. 2D class averaging of a two-chaperonin segment produced a striking similarity to a 2D projection of the crystal structure lattice perpendicular to the “c” axis (Figure 3e). Bundling of filaments, as suggested by the crystal lattice structure of the complex along the “ab” plane, appears to be present when the protein is further enriched resulting in alignment of the striation patterns of adjacent filaments on the micrograph as seen at a protein concentration of 12 mg/mL in the presence of DDM (Supplementary Figure S3, ADP-bound).

Using a single-particle analysis approach in the cryoSPARC2 workflow (Punjani *et al*., 2017), a two-chaperonin segment of the filament could be refined to a resolution of 5.2 Å (Figure 3f, Supplementary Figure S4). Density for the intermediate and apical domains was much clearer than that seen with the chaperonin in isolation, probably as a result of the contacts between the apical domains increasing rigidity this area. Similar to the contacts observed in the crystal structure of TF55β (Supplementary Figure S4b), the protruding helix overlaps with the helix of the adjacent unit, locking each other along the filament (Figure 3g). In the crystal structure, the protruding helices crossed towards the center of the helix. However, MDFF fitting of the cryo-EM model in NAMD (Trabuco *et al*., 2008) minimized on an interaction at the distal end of the helix. This indicated that the interaction resulted primarily from a hydrophobic patch present on the protruding helix (Figure 3h) with a buried surface area of 245 Å^2^ between a pair of protruding helices (Krissinel & Henrick, 2007), which would correspond to ∼2200 Å^2^ between successive octadecamers in a filament. No density attributing to DDM could be observed in the map, though higher resolution studies would likely be needed to observe individual detergent molecules.

### 3.5. Filamentous TF55β form interactions at the protruding helix

The structure of wild-type *S. solfataricus* TF55β filaments indicated that assembly involved an interaction interface in which a hydrophobic patch that is present on the protruding helix becomes buried. This interaction would require the chaperonin to be in an open state for these interfaces to be exposed. Similar interactions between the helical protrusions or adjacent molecules can be seen in the crystal structures of TF55β and the β-only subunit chaperonin from *A. tengchongensis* (Chaston *et al*., 2016; Huo *et al*., 2010). However, the interactions seen in the crystal structures appear to be subtly different to those seen in the filament, involving polar instead of hydrophobic interactions in *S. sulfolobus* and both polar and hydrophobic interactions flanking both sides of the helix in *A. tengchongensis* (Supplementary Figure S4b). Although the interactions seen in the crystals may also be possible in solution, the EM density, albeit at a lower resolution, would suggest that these features are not sufficiently rigid or stable to contribute to the interface seen in the filaments. Such differences in the contacts observed in the crystal structure of TF55β compared to vitrified TF55β single filaments may be ascribed to crystal packing resulting in a helical rise difference of 14 Å (Supplementary Figure S4b). Mutants of *S. shibatae* TF55β have been examined for their ability to form both filaments as well as monolayers that could potentially provide useful materials or coatings for biotechnological applications (Li *et al*., 2007). Surprisingly, removing segments of the protruding helix or the apical domain of TF55β from *S. shibatae* created mutants that were still able to form filaments, possibly suggesting that the overall architecture of the complex, and not purely the helical protrusion, is also capable of assembly into filamentous forms.

Although the function of filament formation in *Sulfolobus* remains obscure, the ability of TF55β to form filaments in solution in the presence of DDM has implications for the production of switchable nanowires.As demonstrated for both TF55β (this study) and GroEL (Chen *et al*., 2015), detergents can induce filament formation and could be used to control this filamentation process in solution on a much shorter timescale than conventional crystallization (Chen *et al*., 2015). High but submicellar concentrations of detergent were required to form detectable and long filaments. NMR studies showed that SDS binds to the inner hydrophobic pocket in their apical domain, which is a known polypeptide substrate-binding site (Saibil *et al*., 2013). TF55β filaments could similarly be disassembled either through dilution (Furutani *et al*., 1998) or through addition of excess EDTA to chelate magnesium and so release nucleotides. Alternatively, as demonstrated for the GroEL-SDS system, cysteine mutants could be designed to rigidify the filament through disulfide bond formation, so that they could also be disassembled using a reducing agent (Chen *et al*., 2015).

## 4. Conclusion

In summary, the cryo-EM structures of TF55 with either ADP or ATP bound show the complex in an open conformation, with an accessible nucleotide binding site that would facilitate nucleotide exchange between chaperoning cycles, whereas our cryo-EM structures of filamentous TF55β highlights the potential interacting surface at the protruding helix during filament assembly.

## Acknowledgements

We wish to thank and acknowledge Jen Coombs (formerly at the School of Biochemistry, University of Bristol) who aided in the initial analysis of this project. We would also like to thank Dr. James Walshe (Victor Chang Cardiac Research Institute) who provided assistance with model building.

## Funding Details

A.G.S was supported by a National Health and Medical Research Council Fellowship APP1159347 and Grant APP1146403. We wish to thank and acknowledge the use of the Victor Chang Cardiac Research Institute Innovation Centre, funded by the NSW Government, and the Electron Microscope Unit at UNSW Sydney, funded in part by the NSW Government.

## Notes

### Competing Interest Statement

The authors have declared no competing interest.

### Summary of Updates

Additional filament formation condition micrographs; Additional dataset of TF55β in complex with ADP showing open structure; Revision of discussion of open TF55β in the presence of nucleotide and filament formation

## References

Adams, P. D., Afonine, P. V., Bunkóczi, G., Chen, V. B., Davis, I. W., Echols, N., Headd, J. J., Hung, L.-W., Kapral, G. J., Grosse-Kunstleve, R. W., McCoy, A. J., Moriarty, N. W., Oeffner, R., Read, R. J., Richardson, D. C., Richardson, J. S., Terwilliger, T. C., Zwart, P. H. & IUCr (2010). Acta Crystallogr. Sect. D Biol. Crystallogr. 66, 213–221.

An, Y. J., Rowland, S. E., Na, J.-H., Spigolon, D., Hong, S. K., Yoon, Y. J., Lee, J.-H., Robb, F. T. & Cha, S.-S. (2017). Nat. Commun. 8, 827.

Bepler, T., Morin, A., Rapp, M., Brasch, J., Shapiro, L., Noble, A. J. & Berger, B. (2019). Nat. Methods. 16, 1153–1160.

Bigotti, M. G. & Clarke, A. R. (2008). Arch. Biochem. Biophys. 474, 331–339.

Chaston, J. J., Smits, C., Aragão, D., Wong, A. S. W., Ahsan, B., Sandin, S., Molugu, S. K., Molugu, S. K., Bernal, R. A., Stock, D. & Stewart, A. G. (2016). Structure. 24, 364–374.

Chaudhry, C., Horwich, A. L., Brunger, A. T. & Adams, P. D. (2004). J. Mol. Biol. 342, 229–245.

Clare, D. K., Vasishtan, D., Stagg, S., Quispe, J., Farr, G. W., Topf, M., Horwich, A. L. & Saibil, H. R. (2012). Cell. 149, 113–123.

Croll, T. I. (2018). Acta Crystallogr. Sect. D Struct. Biol. 74, 519–530.

Emsley, P., Lohkamp, B., Scott, W. G., Cowtan, K. & IUCr (2010). Acta Crystallogr. Sect. D Biol. Crystallogr. 66, 486–501.

Goddard, T. D., Huang, C. C., Meng, E. C., Pettersen, E. F., Couch, G. S., Morris, J. H. & Ferrin, T. E. (2018). Protein Sci. 27, 14–25.

Huo, Y., Hu, Z., Zhang, K., Wang, L., Zhai, Y., Zhou, Q., Lander, G., Zhu, J., He, Y., Pang, X., Xu, W., Bartlam, M., Dong, Z. & Sun, F. (2010). Structure. 18, 1270–1279.

Jonuscheit, M., Martusewitsch, E., Stedman, K. M. & Schleper, C. (2003). Mol. Microbiol. 48, 1241– 1252.

Kagawa, H. K., Yaoi, T., Brocchieri, L., McMillan, R. A., Alton, T. & Trent, J. D. (2003). Mol. Microbiol. 48, 143–156.

Li, Y., Paavola, C. D., Kagawa, H., Chan, S. L. & Trent, J. D. (2007). Nanotechnology. 18, 455101.

Lopez, T., Dalton, K. & Frydman, J. (2015). J. Mol. Biol. 427, 2919–2930.

Muñoz, I. G., Yébenes, H., Zhou, M., Mesa, P., Serna, M., Park, A. Y., Bragado-Nilsson, E., Beloso, A., de Cárcer, G., Malumbres, M., Robinson, C. V, Valpuesta, J. M. & Montoya, G. (2011). Nat. Struct. Mol. Biol. 18, 14–19.

Pereira, J. H., Ralston, C. Y., Douglas, N. R., Kumar, R., Lopez, T., McAndrew, R. P., Knee, K. M., King, J. A., Frydman, J. & Adams, P. D. (2012). EMBO J. 31, 3949–3950.

Pettersen, E. F., Goddard, T. D., Huang, C. C., Couch, G. S., Greenblatt, D. M., Meng, E. C. & Ferrin, T. E. (2004). J. Comput. Chem. 25, 1605–1612.

Pintilie, G. D., Zhang, J., Goddard, T. D., Chiu, W. & Gossard, D. C. (2010). J. Struct. Biol. 170, 427–438.

Punjani, A., Rubinstein, J. L., Fleet, D. J. & Brubaker, M. A. (2017). Nat. Methods. 14, 290–296.

Quehenberger, J., Shen, L., Albers, S.-V., Siebers, B. & Spadiut, O. (2017). Front. Microbiol. 8, 2474.

Saibil, H. (2013). Nat. Rev. Mol. Cell Biol. 14, 630–642.

Saibil, H. R., Fenton, W. A., Clare, D. K. & Horwich, A. L. (2013). J. Mol. Biol. 425, 1476–1487.

Sekiguchi, H., Nakagawa, A., Moriya, K., Makabe, K., Ichiyanagi, K., Nozawa, S., Sato, T., Adachi, S., Kuwajima, K., Yohda, M. & Sasaki, Y. C. (2013). PLoS One. 8, e64176.

Skjærven, L., Cuellar, J., Martinez, A. & Valpuesta, J. M. (2015). FEBS Lett. 589, 2522–2532.

Trent, J. D., Kagawa, H. K. & Yaoi, T. (1998). Ann. N. Y. Acad. Sci. 851, 36–47.

Trent, J. D., Kagawa, H. K., Yaoi, T., Olle, E. & Zaluzec, N. J. (1997). Proc. Natl. Acad. Sci. 94, 5383–5388.

Trent, J. D., Nimmesgern, E., Wall, J. S., Hartl, F.-U. & Horwich, A. L. (1991). Nature. 354, 490– 493.

Trent, J. D., Osipiuk, J. & Pinkau, T. (1990). J. Bacteriol. 172, 1478–1484.

Weiss, C., Jebara, F., Nisemblat, S. & Azem, A. (2016). Front. Mol. Biosci. 3, 80.

Whitelam, S., Rogers, C., Pasqua, A., Paavola, C., Trent, J. & Geissler, P. L. (2009). Nano Lett. 9, 292–297.

Zang, Y., Jin, M., Wang, H., Cui, Z., Kong, L., Liu, C. & Cong, Y. (2016). Nat. Struct. Mol. Biol. 23, 1083–1091.

Zhang, J., Ma, B., DiMaio, F., Douglas, N. R., Joachimiak, L. A., Baker, D., Frydman, J., Levitt, M. & Chiu, W. (2011). Structure. 19, 633–639.

Zivanov, J., Nakane, T., Forsberg, B. O., Kimanius, D., Hagen, W. J., Lindahl, E. & Scheres, S. H. (2018). Elife. 7,.

