## Supporting information for "Structural analysis of *Sulfolobus solfataricus* TF55β chaperonin in open and filamentous states"

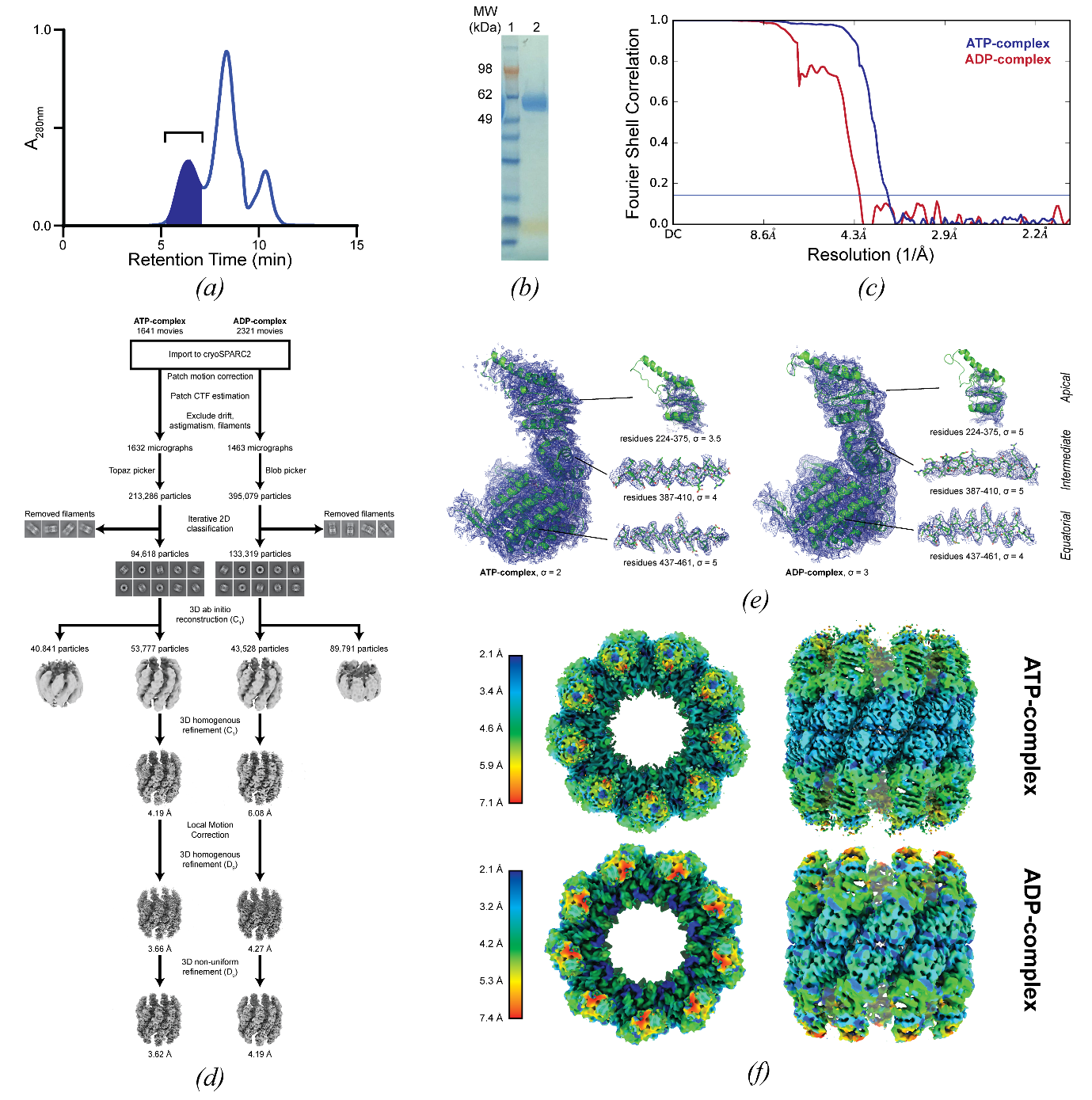

Figure S1 Purification, data processing, and modelling of TF55β chaperonin. *(a)* Representative size exclusion chromatogram after addition of Mg•ATP to purified TF55β monomers. The first peak was collected and concentrated to obtain the TF55β complex. The second highest intensity peak contains TF55β monomers and the third peak contains nucleotides. A similar profile was achieved when Mg•ADP was used. *(b)* SDS-PAGE gel of the TF55β complex after size exclusion chromatograph showing a single protein band corresponding to the TF55β subunit (molecular weight 60 kDa). *(c)* Gold-standard Fourier shell correlation curves (FSC = 0.143, dotted line) of TF55β complex cryo-EM reconstructions showing estimated resolutions of 4.19 Å (red) or 3.62 Å (blue) when 3D refinement was performed for the ADP- and ATP-complex, respectively. *(d)* Flowchart describing cryo-EM data processing for TF55β complex. *(e)* Representative cryo-EM density (blue mesh) for the equatorial, intermediate, or apical domain, contoured at given σ threshold. *(f)* Local resolution estimation performed in cryoSPARC2 for ATP- and ADP-complexes.

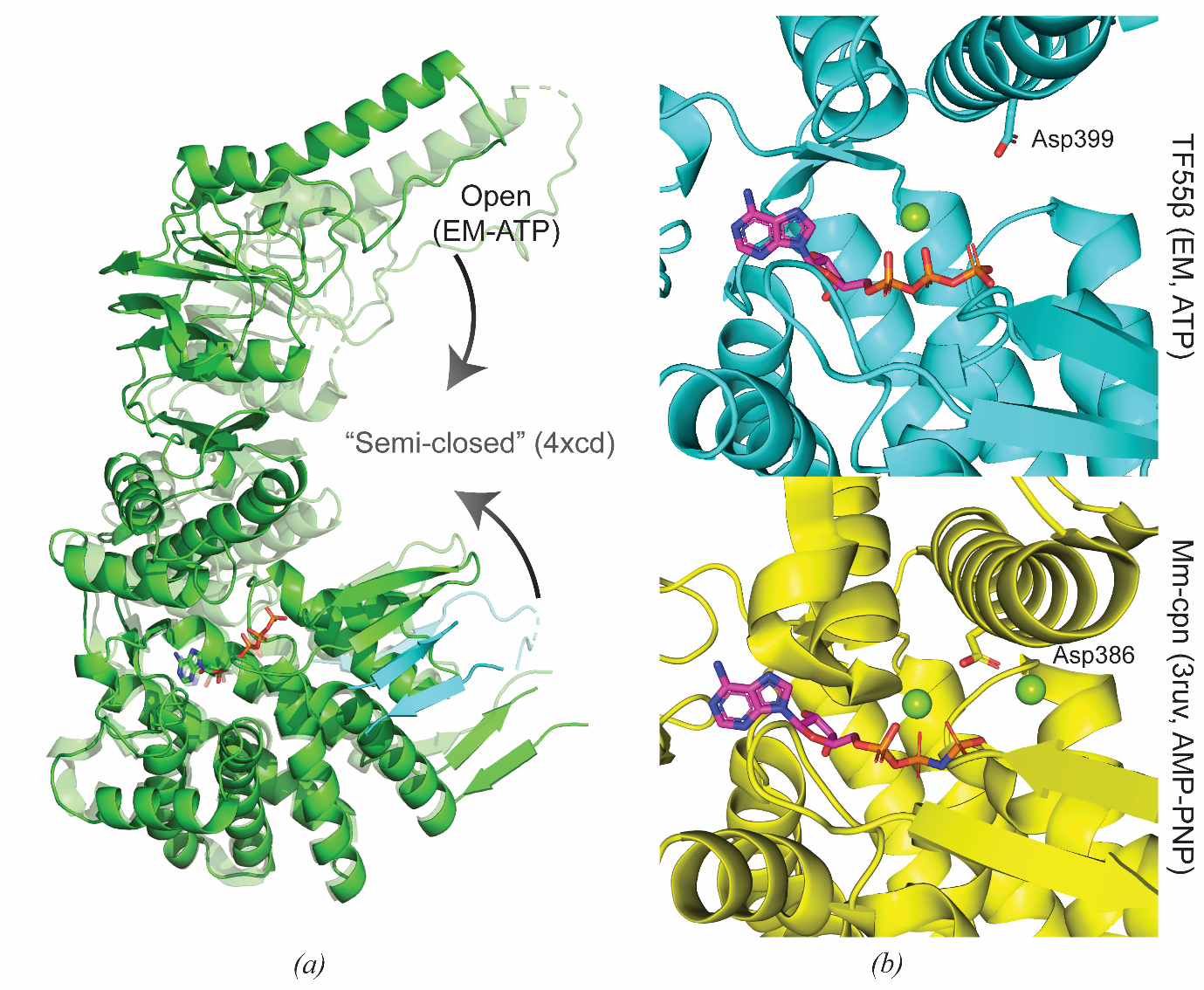

Figure S2 Comparison of open-state TF55β with previous group II chaperonin crystal structures. *(a)* Overlay of cryo-EM structure of TF55β with the “semi-closed” crystal structure (Chaston *et al.*, 2016) showing shifts in the apical domain and the extended β-sheet of the sensor loop/N- and C-termini. Fragment of N- and C- termini β-sheets from the adjacent subunit included in cyan. *(b)* The conserved catalytic aspartate is further away from the magnesium and phosphates of ATP in the open TF55β cryo-EM structure (top) compared to the AMP-PNP-bound structure of *Methanococcus maripaludis* (Pereira *et al.*, 2012) in the closed state (bottom).

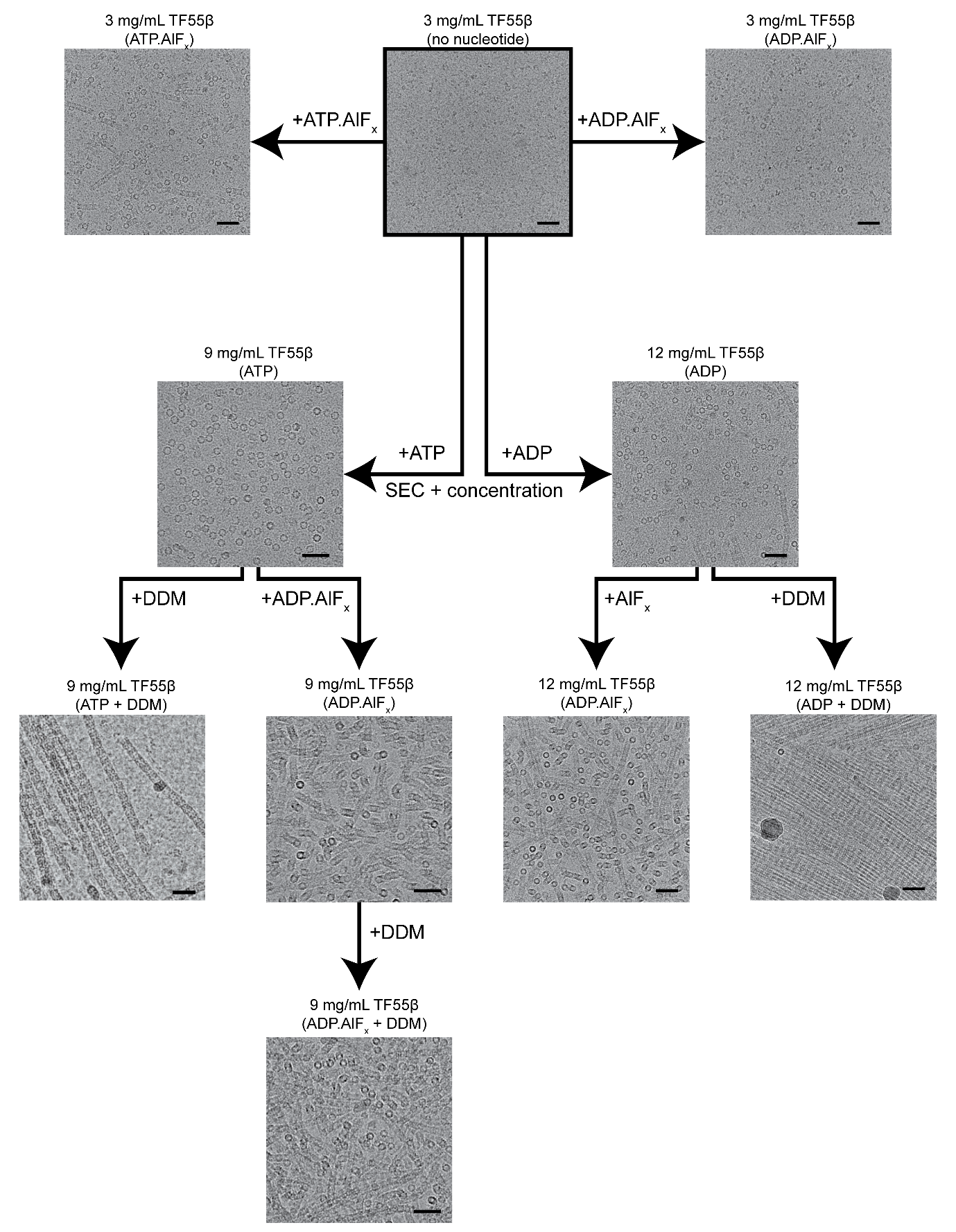

Figure S3 Filament formation of TF55β under different conditions**.** Micrographs of TF55β incubated for 1 h with different reagents to induce filamentation. Scale bars denote 50 nm.

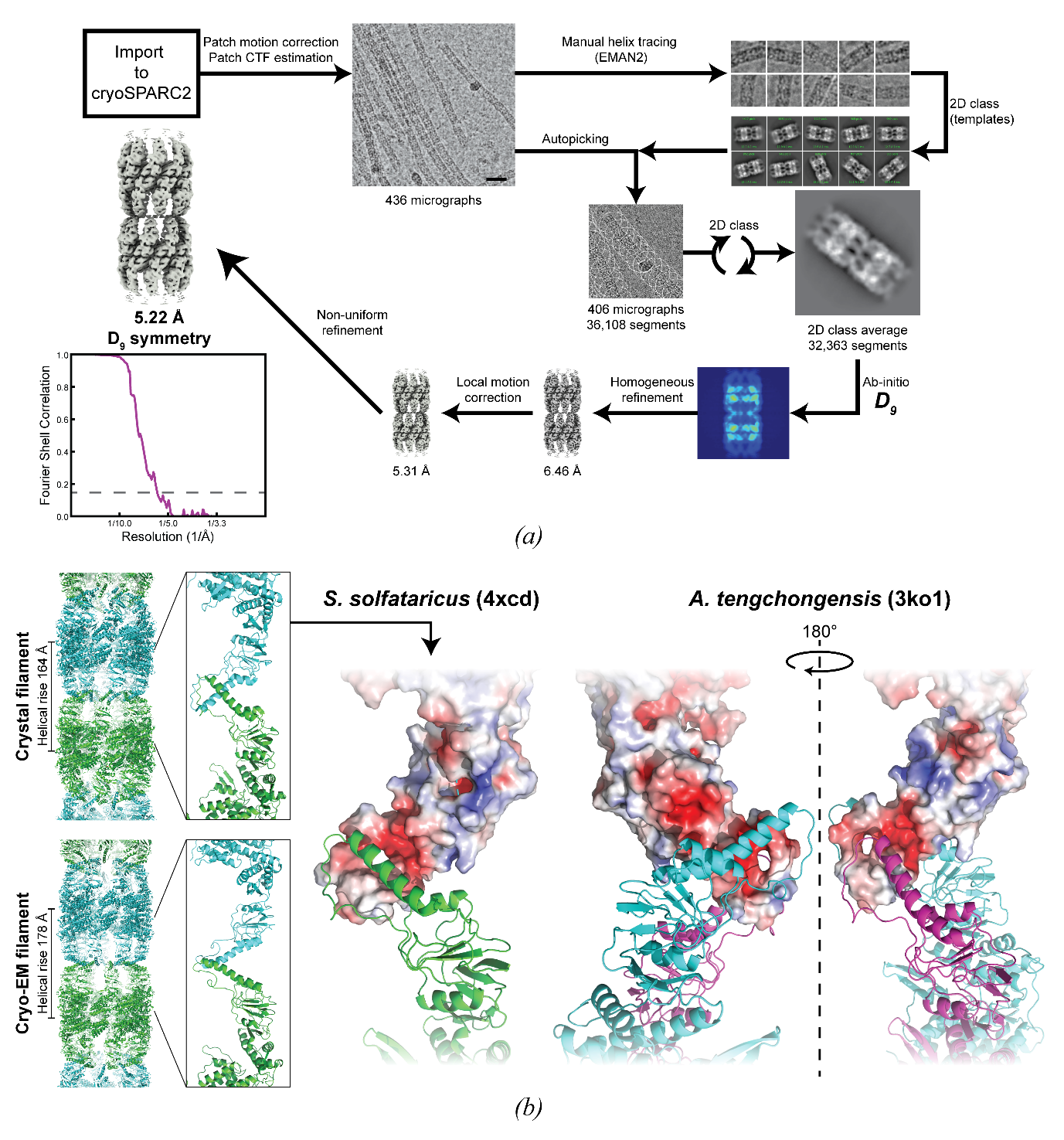

Figure S4 Production of the cryo-EM structure of TF55β filaments. *(a)* Flowchart for obtaining the cryo-EM structure of TF55 filaments. A two-chaperonin segment of filamentous TF55β was processed in cryoSPARC2 to obtain the cryo-EM map with an estimated resolution of 5.22 Å based on the gold-standard Fourier shell correlation criterion (FSC = 0.143, dotted line). *(b)* Comparison of the TF55β filament contacts in the crystal structure (4xcd) and the cryo-EM structure. The protruding helix interfaces at different points. Electrostatic surface representation of the interaction in the crystal structures of TF55β of *S. solfataricus* (Chaston *et al.*, 2016) and *A. tengchongensis* (Huo *et al.*, 2010). *S. solfataricus* TF55β primarily interacts at the polar (red/blue for negative/positive partial charge, respectively) patch of the apical domain, while *A. tengchongensis* apical domain flanks both sides of the helical protrusion at different points.
